# FIQT: a simple, powerful method to accurately estimate effect sizes in genome scans

**DOI:** 10.1101/019299

**Authors:** Tim B. Bigdeli, Donghyung Lee, Brien P. Riley, Vladimir Vladimirov, Ayman H. Fanous, Kenneth S. Kendler, Silviu-Alin Bacanu

## Abstract

Genome scans, including both genome-wide association studies and deep sequencing, continue to discover a growing number of significant association signals for various traits. However, often variants meeting genome-wide significance criteria explain far less of the overall trait variance than “sub-threshold” association signals. To extract these sub-threshold signals, there is a need for methods which accurately estimate the mean of all (normally-distributed) test-statistics from a genome scan (i.e., Z-scores). This is currently achieved by the difficult procedures of adjusting all Z-score 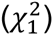 statistics for “winner’s curse” (multiple testing). Given that multiple testing adjustments are much simpler for p-values, we propose a method for estimating Z-scores means by i) first adjusting their p-values for multiple testing and then ii) transforming the adjusted p-values to upper tail Z-scores with the sign of the original statistics. Because a False Discovery Rate (FDR) procedure is used for multiple testing adjustment, we denote this method **F**DR **I**nverse **Q**uantile Transformation (FIQT). When compared to competitors, e.g. Empirical Bayes (including proposed improvements), FIQT is more i) accurate and ii) computationally efficient by orders of magnitude. Its accuracy advantage is substantial at larger sample sizes and/or moderate numbers of association signals. Practical application of FIQT to Z-scores from the first Psychiatric Genetic Consortium (PGC) schizophrenia predicts a non-trivial fraction of the significant signal regions from the subsequent published PGC schizophrenia studies. Finally, we suggest that FIQT might be i) used to improve subject level risk prediction and ii) further improved by modelling the noncentrality of 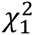 statistics.

## Introduction

Genome-wide association studies (GWAS) represent a powerful and widely used tool for detecting associations between genetic variants and complex traits. In such studies, researchers directly assay and statistically impute (Li, Willer, Ding, Scheet, and Abecasis 2010) genotypes for around a million and several million single nucleotide polymorphisms (SNPs), respectively. The GWAS paradigm has been very successful in identifying genetic variants associated with a range of phenotypes (Dewan, Liu, Hartman, Zhang, Liu, Zhao, Tam, Chan, Lam, Snyder et al. 2006; Hindorff, Sethupathy, Junkins, Ramos, Mehta, Collins, and Manolio 2009; Wellcome Trust Case Control Consortium 2007). However, as seen in GWAS of psychiatric disorders (Purcell, Wray, Stone, Visscher, O’Donovan, Sullivan, and Sklar 2009; Sklar, Ripke, Scott, Andreassen, Cichon, Craddock, Edenberg, Nurnberger, Jr., Rietschel, Blackwood et al. 2011), a considerable portion of the predicted genetic contribution to risk resides in regions which do not independently yield significance at accepted genome-wide levels, i.e., “suggestive” signals. With the advent of large-scale, whole-exome and -genome sequencing studies, the field will likely see an exponential increase in the number of such signals.

Given the increased number such suggestive signals, there is a need for statistical methods that accurately estimate effect-sizes for these and, even, all variants from genome scans (henceforth, denoting not only extant GWAS and whole-exome sequencing but emergent whole-genome sequencing studies as well). These estimates might be used, for instance, to estimate the sample size needed to detect a given number of association signals. Such sample size estimation is, possibly, the most critical consideration in the design of any follow-up sequencing studies.

When estimating the true effect sizes, it is well established that the largest signals are the most affected by the bias known as “winner’s curse” (Zollner and Pritchard 2007). This is due to statistics with the largest magnitude having an extreme value distribution (Jenkinson 1955), as opposed to the normal 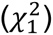 distribution we commonly assume for a random GWAS statistic. By incorrectly assuming a normal distribution, naïve estimators of extreme statistics have a tendency to overestimate the magnitude of these statistics (Zollner and Pritchard 2007). In statistical genetics, researchers proposed a multitude of methods to adjust for winner’s curse studies with one-(discovery) (Faye, Sun, Dimitromanolakis, and Bull 2011; Ghosh, Zou, and Wright 2008; Sun, Dimitromanolakis, Faye, Paterson, Waggott, and Bull 2011; Xiao and Boehnke 2011; Zhong and Prentice 2008; Zollner and Pritchard 2007) and two-stage (discovery and replication) studies (Bowden and Dudbridge 2009; Zhong and Prentice 2008). A majority of these methods are also designed to handle mostly significant signals. Because the statistically significant signals are generally few in number, the majority of these methods are unlikely to meaningfully inform the design of future studies.

Recently, two new tools for estimating all the mean of statistics were proposed. The first was adapted from a more general purpose method from statistics, the Empirical Bayes (EB) method based on Tweedie’s formula (Efron 2009). Because it employs empirical estimates of the density/histogram (120 bins by default) of scan statistics, it is well suited for the large number of statistics from a genome scan (albeit less suited to instances in which the number of statistics is much smaller). In the context of genome scans, this method was used by Ferguson et al. (Ferguson, Cho, Yang, and Zhao 2013), who found that the empirical histogram is less precise in the extreme tails off the distribution, where tail adjustment (TA) methods (Ghosh, Zou, and Wright 2008; Zhong and Prentice 2008) provide better accuracy. Based on these observations, the authors proposed an interesting adaptive combination of EB and TA which, at the cost of increased computational burden, combined the best attributes of both methods. The second of these new tools is a computationally efficient, soft threshold method (Bacanu and Kendler 2012) which adjusts statistics such that their sum of squares do not overestimate the true mean. Because this method does not use empirical density estimation, it is applicable even to a small number of statistics.

We propose a novel, computationally efficient, two-step method to accurately estimate the mean of univariate statistics from a genome scan. First, we adjust the p-values for multiple testing using a False Discovery Rate approach (Benjamini Y. and Hochberg Y. 1995). Second, we estimate the mean of normally distributed statistics as having its i) magnitude equal to the back-transformation of the adjusted p-values to the upper quantiles of a standard normal distribution and ii) sign identical to the original scan statistics. (While more easily explained as a two-step procedure, for the benefit of discussion only, we alternatively present it as a three step procedure in Methods.) As compared to competing methods (with some of our improvements), our proposed procedure has very good performance in terms of i) squared error loss, ii) fraction of the variability in true means of univariate statistics explained and iii) computational efficiency. A practical application of this approach shows that, due to their good performance, the proposed estimators can be used to predict with reasonable accuracy the location of as-yet undetected, significant signals which were ultimately detected in subsequent studies of much larger cohorts. Finally, we suggest that these estimators can be i) possibly used to improve subject level risk prediction and ii) further improved by taking into account the noncentrality of 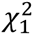 statistics.

## Methods

### Notation

Let *X*_*i*_∼*N*(𝜇_*i*_, 1), *i* = 1,…, *k*, be the univariate statistics from a genome scan and *p*_*i*_, *i* = 1,…, *k*, their associated p-values. If not reported, *X*_*i*_ can be easily computed based on other reported summary statistics (see Supplementary Material). Alternatively, the summary statistics can be reported on a χ^2^ scale with 1 degree of freedom (df), i.e., 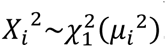. However, in describing the methods we assume the summary statistics are normally distributed (i.e., Z-scores) and their 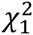 homologs will be used mostly for explaining the heuristics of possible future developments.

### Novel method based on p-value adjustment

Given the extreme value distribution of scan statistics in the upper and lower tails and different distributions elsewhere, it is unclear (or, at least, very complicated) how to properly estimate the biases of, and thereby adjust, all statistics in a genome scan. However, it is quite simple to adjust the p-values for the genome-wide multiple testing, e.g. using a FDR (as in this paper) or Holm procedure. While the FDR procedure can be anticonservative for the extreme scenario of negatively correlated variables, in genetics, Z-scores are only locally correlated and we do not expect their correlations to be preponderantly negative. Thus, FDR is not expected to be conservative and that’s why it is commonly used in statistical genetics. Thus, for a first step let then *p*_*i*_^*^, *i* = 1,…, *k* be the FDR adjusted p-values In the second step, we estimate the expected (adjusted) mean of 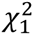 statistics by transforming the adjusted p-values to a 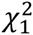 scale 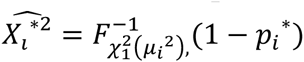 i.e., the upper tail *p*_*i*_^*^ quantile of a 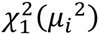 with 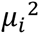 representing the suitably chosen non-centrality parameters used for computing the original p-values *p*_*i*_, *i*= 1,…, *k*. Because the scan statistics are computed under the null hypothesis, H_0_, we used 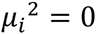 to obtain all of the results presented in this paper (this can be viewed as a shrinkage of *X*_*i*_ toward zero). In the third step, the adjusted standard normal statistic, 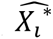 is obtained as 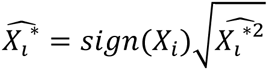. Due to its construction, we denote the proposed method as **F**DR **I**nverse **Q**uantile Transformation (FIQT). Note that, while we have presented FIQT as a three step procedure for the benefit of discussion, it can be simplified to have only two steps. This is achieved by transforming directly on the standard normal scale in the second step, i.e.,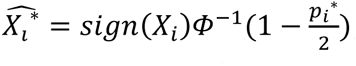 where 𝛷 is the cumulative distribution function (cdf) of a standard normal.

### EB extensions

EB uses all genome scan statistics to i) empirically estimate their density and ii) use the derivatives of the (log) density function to estimate the mean of the statistics and their variance. However, scan statistics are often rather correlated locally (i.e., as a consequence of linkage disequilibrium). This is likely to i) affect the density estimate (which assumes independent statistics) and ii) underestimate the variance of mean statistics. To eliminate (most of) the local correlations we propose an EB extension which i) divides the statistics into *n* equally spaced non-overlapping sets (e.g. first set contains statistics with indices 1, *n* + 1, 2*n* + 2,…, and the second those with indices 2, *n* + 2, 2*n* + 3,…,), ii) estimates the density for each set, iii) uses each set density to estimate a set-specific mean for all scan statistics and iv) estimates the overall mean of scan statistics as the average set-specific means. We denote this estimator as EB-*n*, i.e., when using 100 non-overlapping sets the EB extension is denoted as EB-100. The obvious disadvantage of EB-*n* over EB is its increased computational burden, as the computationally intensive estimation of density and its derivatives are computed *n* times.

### Methods used for comparisons

For comparison, we use the naïve maximum likelihood estimator (MLE), i.e., the statistics themselves, classical EB (EB-1 in the above set notation) and EB-*n* (*n*=10, 50, 100). (Because the soft threshold method (Bacanu and Kendler 2012) was found to slightly underperform EB-1, for brevity, we omit it from our results). Due to it sometimes outperforming EB in the tails, we also include the tail adjustment (TA) method (Ghosh, Zou, and Wright 2008; Zhong and Prentice 2008). The original TA adjusts all statistics above a preset (and generally significant) threshold, which results in two unusual features for our presentation of results. First, given that remaining methods adjust all statistics in a genome scan, we employed TA outside its intended purposes, e.g. even for (very) non-significant thresholds. Second, given TA’s approach of computing the bias for all statistics above a signal threshold, we present the performance of tested methods (MSE and R^2^ in Results section) in a cumulative manner, i.e. for all statistics with p-values below a large range of thresholds.

### Implementation and assessment of performance

We implemented all described methods using the R statistical programming environment. For FDR, FIQT employed the *p.adjust* base function with the *“fdr“* option specified for method (see SM for the 10-line implementation in R). EB methods employed the commonly used 120 bins for the empirical estimation of the probability density. The running time of tested methods was assessed using the second entry (i.e. *“system“*) in the output of *system.time* function.

### Simulations

Simulating genome scan statistics starting from genotypes is laborious and very time consuming. Consequently, we use a faster method that simulates scan statistics directly using an ARMA (3,4) model for residuals [see Simulation model in Supplementary Material (SM) and Table I]. This model was found to be adequate for simulating statistics for markers with a density of approximately 1 SNP/kbp (Bacanu and Kendler 2012). Effect sizes (i.e. the true mean of the statistics) and number of the signals were based on their homologs in a mega-analysis of human height (Lange, van, Andrew, Lyon, DeMeo, Raby, Murphy, Silverman, MacGregor, Weiss et al. 2004). We assumed that the phenotype under investigation has *m*_1_ causal loci which represent a fraction *γ*_*c*_ ≤ 1 of the number of significant loci (*m* = 180) in height study (Table I), i.e.*m*_1_ = *γ*_*c*_ *m*. When *γ*_*c*_ < 1, the *m*_1_ causal loci are chosen at random from the significant loci in the height study. To assess the performance of methods for underpowered studies, we performed simulations under *H*_0_. Under this scenario *γ*_*c*_ = 0, i.e. the simulated statistics are identical to an ARMA (3,4) realization of unit variance. We simulated sample sizes equaling a fraction 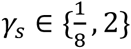 of the height meta-analysis sample size (*n* ≈ 180,000). Additional details regarding the relationship between mean of the statistics and *γ*_*s*_, are available in the Simulation model subsection in SM. For every parameterization given in Table I, we performed 250 simulations.

**Table I.** Simulation design parameters.

| Parameter Name | Parameter | Design levels |
| --- | --- | --- |
| ARMA(3,4) model used for simulating statistics | AR vector | $\{0.8716, 0.9782, -0.851\}$ |
| | MA vector | $\{-0.6652, -0.9976, 0.6594, 0.0252\}$ |
| Number of simulated autosome SNPs | $k$ | 2,866,105 (1 SNP/Kbp) |
| Phenotyped sample size (thousands) | $n_1$ | $\{22.5, 45, 90, 180, 360\}$ (i.e. fraction of height meta-analysis sample $\gamma_s = \{\frac{1}{8}, \frac{1}{4}, \frac{1}{2}, 1, 2\}$) |
| Number of causal SNPs | $m_1$ | $\{6, 11, 23, 45, 90, 180\}$ (i.e. fraction of height meta-analysis number of causal SNPs $\gamma_c = \{0, \frac{1}{32}, \frac{1}{16}, \frac{1}{8}, \frac{1}{4}, \frac{1}{2}, 1\}$) |

### Practical Application

We applied FIQT to the discovery phase of the 2005 Psychiatric Genomics Consortium (PGC) GWAS of schizophrenia (PGC-SCZ1) (Ripke, Sanders, Kendler, Levinson, Sklar, Holmans, Lin, Duan, Ophoff, Andreassen et al. 2011) to naïvely estimate which genomic regions harbored statistics expected to attain significance in four-fold larger discovery phase of the 2014 PGC study (PGC-SCZ2) (Schizophrenia Working Group of the Psychiatric Genomics Consortium 2014).

### Results

Among EB-*n* methods, we tested EB-1, EB-10, EB-50 and EB-100. EB-10 and EB-50 have intermediate performance between EB-1 and EB-100, with EB-10 closer to EB-1 and EB-50 closer to EB-100 (data not shown). Consequently, we present only the results for EB-1 and EB-100.

Under *H*_0_, i.e. the surrogate for underpowered studies, FIQT has the best mean square error (MSE) performance everywhere, except for the (very rare) region of extremely low p-values, where EB-1 slightly outperforms it (Fig.1). Among the remaining methods, EB-1 performs best and, as expected, MLE has the largest MSE. We note that, in marked contrast to the alternative hypothesis results presented below, EB-1 thoroughly outperforms EB-100.

**Figure 1.**
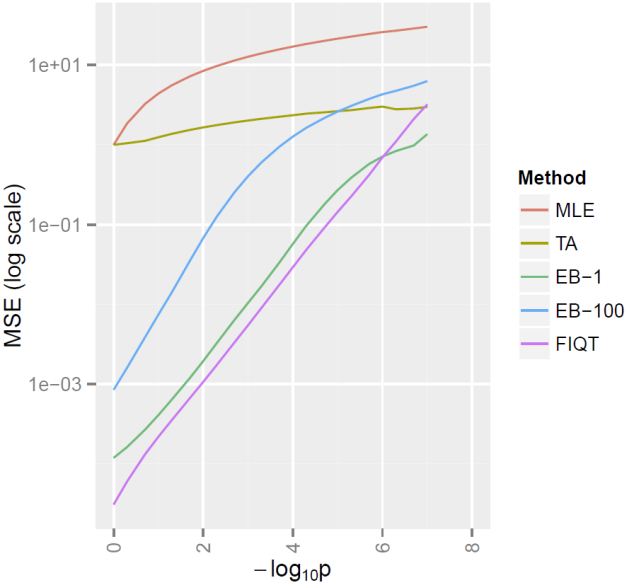
Null hypothesis mean square error (MSE) of FIQT estimates for Z-scores having negative log p-values below a threshold equal to −log_10_p.

Under the alternative, *H*_*a*_, FIQT has better MSE performance for settings with larger number of signals and larger sample sizes (Fig.2). Its performance improvement over competitors is sometimes substantial, e.g. for large sample sizes and medium number of signals. EB-1 does not outperform FIQT under any *H*_*a*_ scenarios and EB-100 only nominally so for smaller sample sizes. Surprisingly, even though it was designed only as a tail bias adjustment, TA performs reasonably well. Under certain scenarios, e.g. large sample sizes, it outperforms EBs for statistics with nominally significant p-values and even slightly outperforms FIQT for a very narrow range of moderately small p-values. When the accuracy measure is R^2^, i.e. the explained variability of the true Z-scores means, FIQT practically outperforms all other methods (Fig. 3), albeit EB-100 only nominally.

**Figure2.**
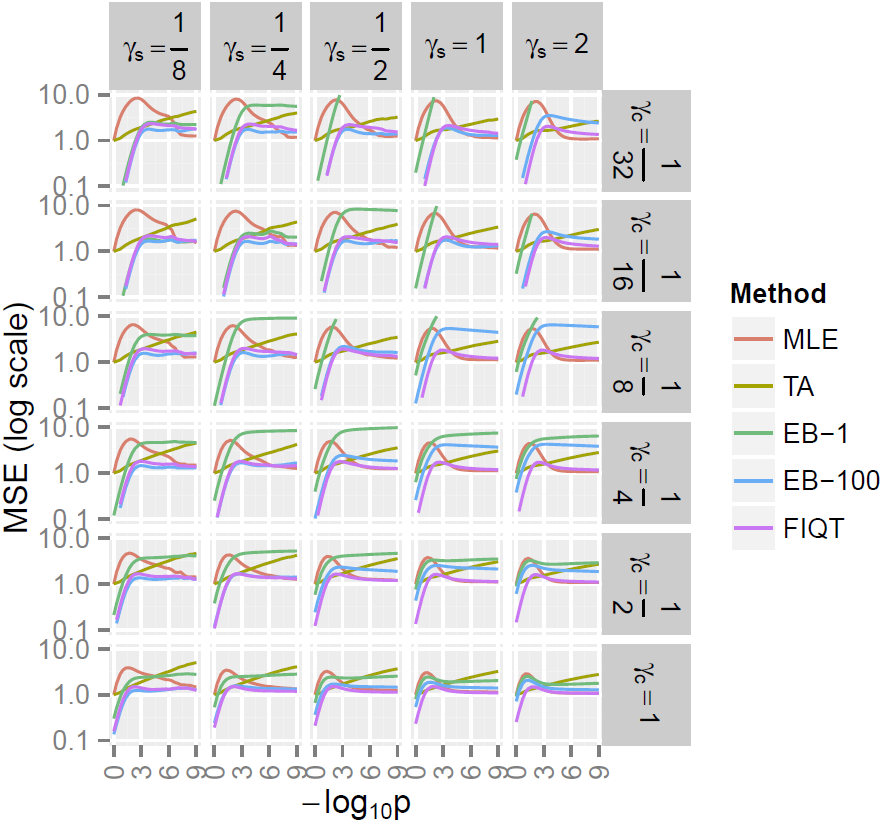
Alternative hypothesis FIQT MSE for Z-scores having negative log p-values below −log_10_p. *γ*_*s*_ is the relative (to height meta-analysis) sample size and *γ*_*c*_ is the number of causal signals relative to 180 significant height meta-analysis signals.

**Figure 3.**
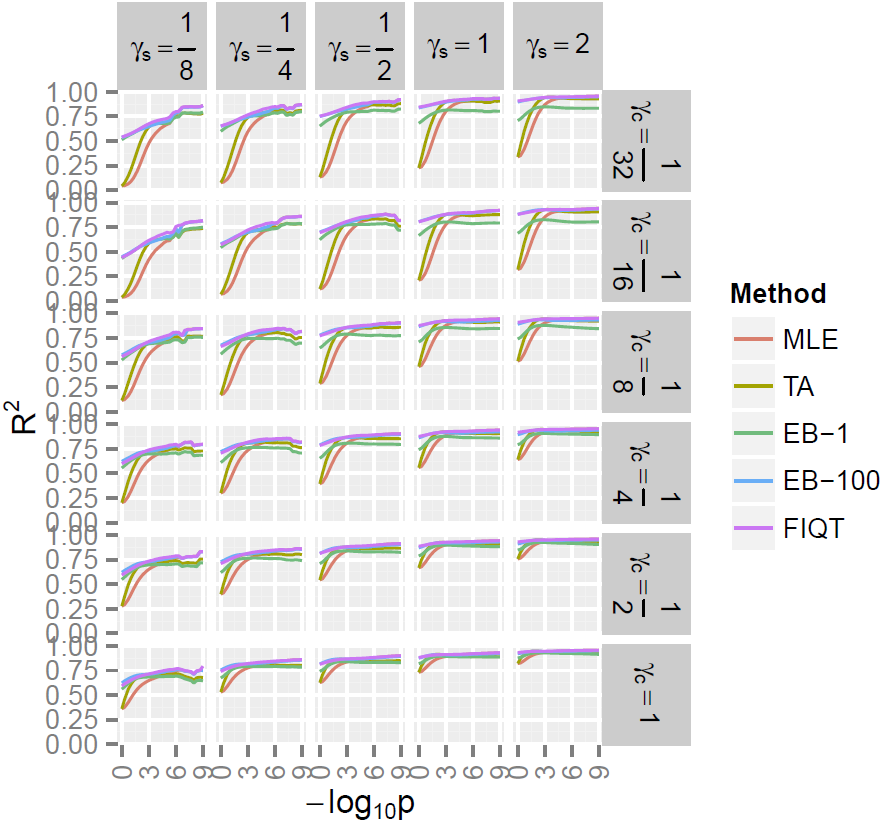
The alternative hypothesis variability in true means (R^2^) for Z-scores having negative log p-values below −log_10_p. See Fig.2 for background and notations.

Due to its very simple computation, FIQT has much faster running times then competitors. When compared to the next most accurate method, EB-100, the proposed method is faster by more than four orders of magnitude (Fig. S1 in SM). FIQT is also faster than the less accurate EB-1 by almost two orders of magnitude (data not shown).

### Practical application

Given that discovery phase of PGC-SCZ2 has around four times the sample size of its PGC-SCZ1 homolog, then naïvely, the mean statistics in PGC2 are expected to be twice as large as the PGC-SCZ1 FIQT estimates (see Fig. S2 for relationships between these estimators and PGC-SCZ1 statistics). This naïve FIQT application to PGC-SCZ1 resulted in the prediction of 46 regions expected to attain significance in PGC-SCZ2 (obtained by combining significant signals within 250 Kb). Of these, 34 regions (∼ 75%) overlap the 105 independent chromosomal regions reported in PGC-SCZ2. A total of 18 predicted PGC-SCZ2 regions overlap the extended MHC region (25-33 Mb on chromosome 6) from the actual PGC-SCZ2 findings, as opposed to only 5 reported by PGC-SCZ1. Of the 34 overlapping predicted PGC-SCZ2 regions, 16 are in loci outside MHC regions, as opposed to just 4 reported by PGC-SCZ1. These non-MHC regions include *CACNA1C* and the *ITIH3/ITIH4* cluster, which were reported as harboring significant signals only after jointly analyzing PGC-SCZ1 and bipolar disorder cohorts (Ripke, Sanders, Kendler, Levinson, Sklar, Holmans, Lin, Duan, Ophoff, Andreassen et al. 2011).

## Discussion

We propose a novel approach, FIQT, to extract information from genome scan statistics by accurately predicting the mean of the statistics. The accurate prediction is achieved via a three-step process. First, the p-values are FDR adjusted for multiple testing. Second, we estimate the square of mean statistics by transforming the adjusted p-values to a upper 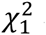 quantiles. Third, we estimate the mean of normally distributed scan statistics as the square root of the adjusted 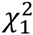 quantile with the sign of the unadjusted Z-score statistics. (The second and third steps can be combined into a single step by transforming to a normal quantile.) When compared to competing methods, FIQT estimators are shown to have a i) smaller mean squared error loss, ii) explain a higher proportion of the true means of the statistics and iii) substantially faster running times. The practical application to PGC-SCZ1 summary statistics show that FIQT estimators are useful for highlighting, with reasonable specificity, genomic regions likely to show significant signals in (much larger) future studies.

Empirical Bayes, EB, and similar methods are currently some of the state-of-the-art approaches for accurately estimating the true means of scan statistics. However, in its commonly used form, EB does not take into account the local LD dependence between SNPs. We proposed EB-*n*, which eliminate this dependence by subdividing the SNPs in *n* non-overlapping sets. These methods, e.g. EB-100, improve over the performance of classical EB, albeit at the expense of a dramatically increased computation time. However, all EB methods are computationally and skill intensive and, due to the need of empirically estimating the probability density of statistics, they are not applicable when the number of statistics is reasonably small. Our proposed method, FIQT, eliminates these disadvantages while maintaining a similar or (sometimes much) better prediction accuracy. Its performance advantage over EB based methods is substantial at high sample sizes and a moderate to large number of true signals.

FIQT is designed to estimate the true means for all Z-scores in the genome scan. Sometimes, Z-scores are not present and the researchers need to estimate them from other summary data (see SM). Conversely, the adjusted Z-scores (e.g. FIQT estimates of their true mean) can be subsequently used to estimate the adjusted vales for the original (reported) summary statistics. For instance, if summary data contain only odds ratio, OR, and their standard error, s, then the vector of Z-scores is 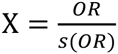. Subsequently, we can use the adjusted Z-scores (FIQT estimates), X*, to estimate the adjusted odds ratio, e.g. as OR* = sX* (or, simply 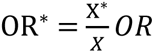). OR* can be interpreted as the vector of unbiased (winner’s curse corrected) ORs.

FIQT is a very simple yet powerful method. However, in its present form, it is more of a proof-of-concept and it can be further improved. A first direction of improvement is to shrink the 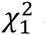 statistics towards their actual mean, not towards zero as used in our simulations and practical application. A second improvement might involve shrinking the 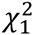 statistics towards their functional group mean, i.e. somewhat similar to the conditional (p-value based) approach from Andressen et al (Andreassen, Djurovic, Thompson, Schork, Kendler, O’Donovan, Rujescu, Werge, van de Bunt, Morris et al. 2013). A second possible improvement might be realized by taking into account the correlation between statistics.

FIQT can also be extended to the accurate estimation of other variables besides the mean of Z-scores. For instance, shrinkage estimators are widely used for correlation/covariance matrices (Daniels and Kass 2001). Given that the sample correlations are normally distributed with variances dependent only on the sample size, FIQT can be extended to the estimation of correlation matrices. The extension might involve shrinking the magnitudes of entries in the correlation matrix toward i) zero, ii) their average magnitude or iii) their average magnitude by lag.

FIQT might be also used in the personalized genomics, e.g. the prediction of subject level risk based on whole genome data. Methods for predicting subject level risk typically use summary statistics as input, e.g. LDpred extension (http://biorxiv.org/content/early/2015/03/04/015859) of LD score method (Bulik-Sullivan, Loh, Finucane, Ripke, Yang, Patterson, Daly, Price, and Neale 2015). Thus an increased accuracy of signal estimation used as input will result in more accurate estimates of an individual’s risk/trait mean.

### Software

A robustly tested FIQT R function having only 10 lines of code is available in the third section of SM. FIQT will also be implemented in DIST-MIX, or our group’s imputation software for cosmopolitan cohorts (https://code.google.com/p/distmix/).

## Acknowledgements

This work was supported by R25DA026119 (D.L.), MH100560 (S.A.B. and B.P.R.), AA022717 (S.A.B.) and 1P50AA022537 (S.A.B. and B.P.R.).

